# Generation of Variation and Mean Fitness Increase: Necessity is the Mother of Genetic Invention

**DOI:** 10.1101/229351

**Authors:** Yoav Ram, Lee Altenberg, Uri Liberman, Marcus W. Feldman

**Affiliations:** Department of Biology, Stanford University, Stanford, CA; Information and Computer Sciences, University of Hawai‘i at Mānoa, Honolulu, HI; School of Mathematical Sciences, Tel Aviv University, Israel

## Abstract

Generation of variation may be detrimental in well-adapted populations evolving under constant selection. In a constant environment, genetic modifiers that reduce the rate at which variation is generated by processes such as mutation and migration, succeed. However, departures from this *reduction principle* have been demonstrated. Here we analyze a general model of evolution under constant selection where the rate at which variation is generated depends on the individual. We find that if a modifier allele increases the rate at which individuals of below-average fitness generate variation, then it will increase in frequency and increase the population mean fitness. This principle applies to phenomena such as stress-induced mutagenesis and condition-dependent dispersal, and exemplifies *“Necessity is the mother of genetic invention.”*

## Introduction

According to the *reduction principle*, in populations at a balance between natural selection and a process that generates variation (i.e. mutation, migration, or recombination), selection favors neutral modifiers that decrease the rate at which variation is generated. The *reduction principle* was demonstrated for modifiers of recombination (Feldman, 1972), mutation (Liberman and Feldman, 1986), and migration (Feldman and Liberman, 1986). These results were unified in a series of studies (Altenberg, 1984; Altenberg and Feldman, 1987; Altenberg, 2009, 2012a,b; Altenberg et al., 2017).

The latter studies have established the conditions for a *unified reduction principle* by neutral genetic modifiers: (i) effectively infinite population size, (ii) constant-viability selection, (iii) a population at an equilibrium, and (iv) *linear variation* – the equal scaling of transition probabilities by the modifier. A departure from the latter assumption occurs if two variation-producing processes interact (Feldman et al., 1980; Altenberg, 2012a). Departures from the *reduction principle* have also been demonstrated when conditions (i)-(iii) are not met, see for example Holsinger et al. (1986) and references therein.

Another departure from the *linear variation* assumption of the *reduction principle* for mutation rates involves a mechanism by which the mutation rate increases in individuals of low fitness – a mechanism first observed in stressed bacteria (Foster, 2007), although not in a constant environment. Ram and Hadany (2012) demonstrated that even in a constant environment, increasing the mutation rate of individuals with below-average fitness increases the population mean fitness, rather than decreases it. Their analysis assumed infinite population size and fitness determined by the number of mutant alleles accumulated in the genotype. In their models, the only departure from the *reduction principle* assumptions was the unequal scaling of mutation probabilities between different genotypes introduced by the correlation between the mutation rate and fitness. A similar result has been demonstrated for conditional dispersal (Altenberg, 2012a, Th. 39), fitness-associated recombination (Hadany and Beker, 2003*b*) and for condition-dependent sexual reproduction (Hadany and Otto, 2007), and evidence suggests that both mechanisms are common in nature (Ram and Hadany, 2016).

Ram and Hadany (2012) stated that their result represents a departure from the *reduction principle*, but did not explain this departure. Their analysis was specific to a model that classified individuals by the number of mutant alleles in their genotype, similar to models studied by Kimura and Maruyama (1966) and Haigh (1978). Moreover, their argument was based on the expected increase of the stable population mean fitness, rather than on the invasion success of modifier alleles that modify the mutation rate (i.e., analysis of *evolutionary genetic stability,* see Eshel and Feldman, 1982; Lessard, 1990).

Here, we present an evolutionary model in which the type of the individual determines both its fitness and the rate at which it generates variation. Our results show that the population mean fitness increases if individuals with below-average fitness produce more variation than individuals with above-average fitness, and that modifier alleles that induce below-average individuals to produce more variation are favored by natural selection.

## Models

### General model

Consider a large population with an arbitrary set of types *A*_1_, *A*_2_,…, *A*_*n*_. The frequency and fitness of individuals of type *A*_*k*_ are *f_k_* and *W_k_*, respectively. The probability that an individual of type *A*_*k*_ will transition to some other type is *C_k_*, and given a transition occurs, the probability that it will transition to type *A_j_* is *M_j,k_*. Therefore, the change in the frequencies of type *A_k_* is described by the transformation *f* → *f′*:

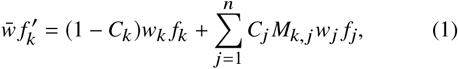

or in matrix form

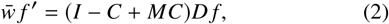

where *f* = (*f*_1_, *f*_2_,…, *f_n_*) is a frequency vector with *f*_*k*_ ≥ 0 and 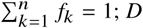 is a positive diagonal matrix with entries *w_k_* such that *w_k_* ≠ *w_j_* for some *k* ≠ *j*; *C* is a positive diagonal matrix with entries *C_k_*; *M* is a primitive column-stochastic matrix: *M_j,k_* ≥ 0 for all 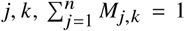 for all *k*, and (*M^l^*)_*j,k*_ > for all *j*, *k* for some positive integer *l*; *I* is the *n* × *n* identity matrix; and *w̅* is the normalizing factor such that 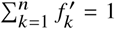 and is equal to the population mean fitness 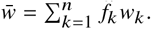

The types *A_k_* can represent a single or multiple haploid genetic loci or non-genetic traits. Importantly, type transmission is vertical and uni-parental (the type is transmitted from a single parent to the offspring) and is independent of the frequencies of the other types. This model precludes processes such as recombination, social learning, sexual outcrossing, and horizontal or oblique transmission, as these processes are frequency-dependent (Cavalli-Sforza and Feldman, 1981, pg. 54).

Transition between types is determined by a combination of two effects: (i) the probability of transitioning *out* of type *A_k_* is determined by *C_k_*; (ii) given a transition out of type *A_k_*, the distribution of the destination types *A_i_*, is given by *M_i,k_* (note the index order). Importantly, different types can have different rates. That is, *C_i_* ≠ *C_j_* for some *i, j*. The case *C_i_* = *C_j_* for all *i, j* is covered by the *reduction principle* (see Altenberg et al., 2017).

In the following section we present four examples of the model (Eq. 2) that apply to mutation, migration, and learning.

### Mutation model 1

Here we consider a large population of haploids and a trait determined by a single genetic locus with *n* possible alleles *A*_1_, *A*_2_, …, *A_n_* and corresponding fitness values *w*_1_, *w*_2_, …, *w_n_*. The mutation rates *C_k_* of individuals with allele *A_k_* are potentially different; specifically, with probability 1 – *C_k_*, the allele *A_k_* does not mutate, and with probability 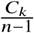, the allele *A_k_* mutates to *A_j_* for any *j* ≠ *k*. This is an extension of a model studied by Altenberg et al. (2017) that allows for the mutation rate of *A_k_*, *C_k_*, to depend on properties of the allele *A_k_*.

Let the frequency of *A_k_* in the present generation be *f_k_* with *f_k_* ≥ 0 and 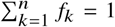. Then after selection and mutation, 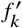 in the next generation is given by

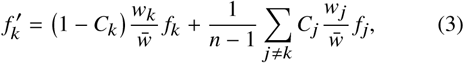

for *k* = 1, 2,…, *n*, where 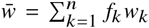 is the population mean fitness.

This model is a special case of the general model (Eq. 2) where

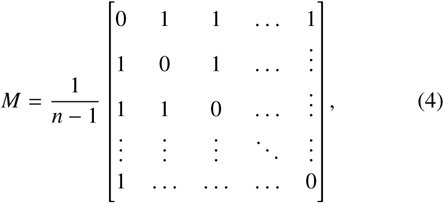

with zeros on the diagonal and 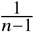 elsewhere. Note that here *M* is irreducible and primitive.

### Mutation model 2

Again, we consider a large population of haploids, but here individuals with genotype *A_k_* are characterized by the number *k* of deleterious or mutant alleles in their genotype, where 0 ≤ *k* ≤ *n*. Specifically, the fitness of individuals with *k* mutant alleles is *W_k_* (*W*_0_ > *W*_1_ > … > *w_n_*), and the probability *C_k_* that a mutation occurs in individuals with *k* mutant alleles depends on *k*. When a mutation occurs it is *deleterious* with probability *δ*, generating a mutant allele and converting the individual from *A_k_* to *A*_*k*+1_, or it is *beneficial* with probability *β*, converting the individual from *A_k_* to *A*_*k*−1_. Note that such beneficial mutations can be either compensatory or back-mutations, and that mutations are neutral with probability 1 – *δ* – *β*. We assume that both the deleterious and the beneficial mutation rates are low enough that two mutations are unlikely to occur in the same individual in one generation: *C_k_* (*δ* + *β*) ≪ 1 for all *k* = 1,…, *n*. This model has been analyzed by Ram and Hadany (2012).

Let the frequency of *A_k_* in the present generation be *f_k_* with *f_k_* ≥ 0 and 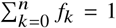. Then after selection and mutation 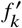 in the next generation is given by

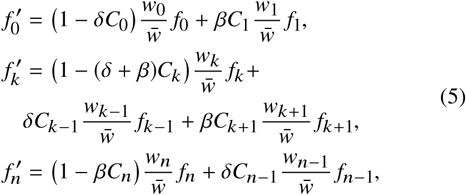

for *k* = 1, 2,…, *n* – 1. Here 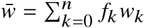 is the population mean fitness.

Therefore, setting

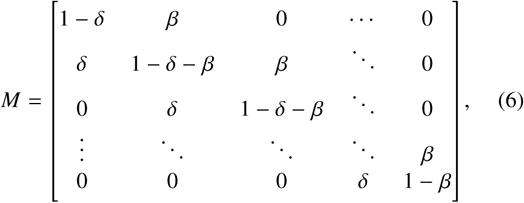

with the beneficial, neutral, and deleterious mutation probabilities on the three main diagonals and zeros elsewhere, Eq. 5 can be viewed as a special case of Eq. 2. Here, too, *M* is irreducible and primitive as long as *δ, β* > 0.

### Migration model

In this case we consider a large population of haploids that occupy *n* demes, *A*_1_,…, *A_n_*. Let the frequencies of individuals in deme *A_k_* be *f_k_* with *f_k_* ≥ 0 and 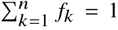. The fitness of individuals in deme *A_k_* is *w_k_*, but the entire population comes together for reproduction, and therefore reproductive success is determined by competition among individuals of all demes – this has been termed *hard selection* (Wallace, 1975; Karlin, 1982).

After reproduction, offspring of individuals from deme *A_k_* return to their parental deme with probability 1 – *C_k_*, or migrate to a different deme *A_j_* with probability *C_k_M_j,k_*, where the matrix *M* is primitive, *C_k_* > 0, *M_j,k_* ≥ 0, and 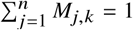 for all *k* = 1,…, *n*. Therefore, 1 – *C_k_* can be interpreted as a *homing rate.*

Following selection and migration the new frequencies 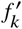 are given exactly by Eq. 1. If the columns of *M* are identical

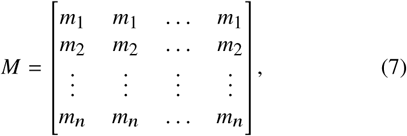

with *m_k_* > 0 and 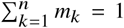, then *m_k_* can be considered the relative population size of deme *A_k_* – this is the non-homogeneous extension of Deakin’s *homing model* (Deakin, 1966; Karlin, 1982).

Similarly, if demes are arranged in a circle, for example around a lake, then we can denote the probability *p_k_* of migrating *k* demes away from the parental deme (conditioned on migration which occurs with probability *C_k_*) and *M* takes the form

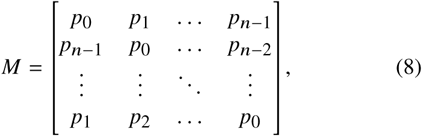

where *p_k_* > 0 and 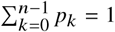.

### Learning model

In our final example, we consider a large population and an integer phenotype *k* where 1 ≤ *k* ≤ *n*. Individuals are characterized by their initial and mature phenotypes (Boyd and Richerson, 1985, pg. 94). Fitness is determined by the mature phenotype: the fitness of an individual with mature phenotype *k* is *w_k_*.

An offspring’s initial phenotype is acquired by learning the mature phenotype of its parent (assuming uni-parental transmission). In individuals with initial phenotype *k*, the mature phenotype is the same as the initial phenotype with probability 1 – *C_k_*, and is modified by individual learning or *exploration* (Borenstein et al., 2008) with probability *C_k_*. Such individual exploratory learning, which can be considered either intentional or the result of incorrect learning, modifies initial phenotype *k* to mature phenotype *j* with probability *M_j,k_*.

Therefore, if the frequency of individuals with mature phenotype *k* in the current generation is *f_k_*, then the frequency in the next generation 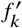 is

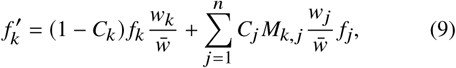

where 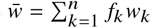 is the population mean fitness.

For example, in the case of *symmetric individual learning* (Borenstein et al., 2008), learning is parameterized by its breadth of exploration *b* and the mature phenotype *j* is randomly drawn from 2*b* + 1 phenotypes symmetrically and uniformly distributed around the initial phenotype *k*, with the limitation that any “spillover” of phenotypes below 1 or above *n* is “absorbed” by those boundaries. This “absorption” ensures *M* is column-stochastic. In other words, given initial phenotype *k*, the probability of maturation to phenotype *j*, where *k* – *b* ≤ *j* ≤ *k* + *b* is 1/(2*b* + 1), but any phenotype *j* < 1 actually becomes *j* = 1 and any phenotype *j* > *n* actually becomes *j* = *n*. The probability for maturation to other phenotypes is 0.

For instance, with *n* = 5 and *b* = 1 we have

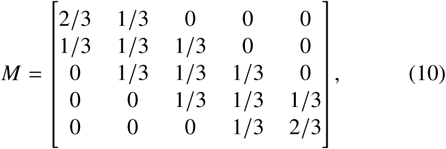

and *n* = 6 and *b* = 2 we have

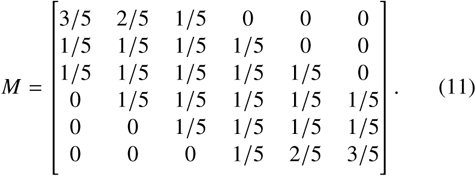

## Results

### Mean fitness principle

We first focus on the stable population mean fitness. We show that if the transition rate from types with below-average fitness increases, then the stable population mean fitness increases, too.

Write the equilibrium frequency vector *f* in Eq. 2 as 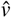 and the stable population mean fitness as 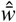, then

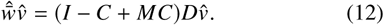

Note that (i) the existence and uniqueness of 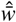 and 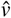 are guaranteed by the *Perron-Frobenius theorem* (Otto and Day (2007)) because (*I* – *C* + *MC*)*D* is a non-negative primitive matrix; (ii) the global stability of this equilibrium is proven in Appendix C.

The following result constitutes a *mean fitness principle* for the sensitivity of the equilibrium mean fitness 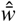 to changes in *C_k_*, the probability of transition from *A_k_*.

### Result 1 (Mean fitness principle)

*Let 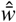 be the leading eigenvalue of* (*I* – *C* + *MC*)*D*, *and û and* 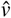 *be the corresponding positive left and right eigenvectors, such that* 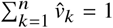 *and* 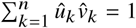. *Then,*

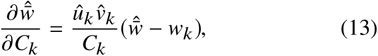

*or in simpler terms,*

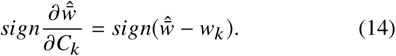

*Therefore increased transition from type k will increase the stable population mean fitness if the fitness of type k is below the stable population mean fitness.*

#### Proof

Using the formula in Caswell (1978) (see Eq. 36 in Appendix A),

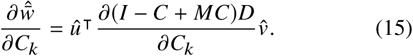

Let *e_k_* and 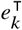 be the column and row vectors with 1 at position *k* and 0 elsewhere, 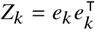 be the matrix with 1 at position (*k, k*) and 0 elsewhere, and [*M*]*k* be the *k*-th column of *M*.

Then,

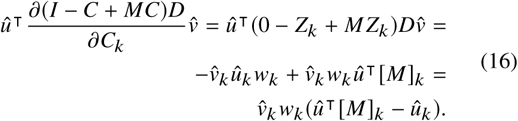

The corresponding equation to Eq. 12 for the left eigenvector *u* is

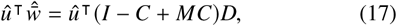

which gives us a relation between 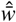 and the *k* element of *û*:

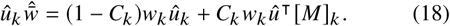

Multiplying both sides by 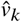 and rearranging, we get

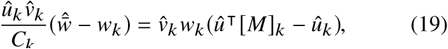

which when substituted into Eq. 16 yields:

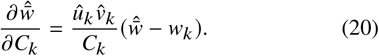

Finally, since 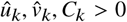, we have

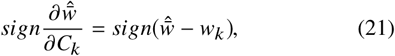

which completes the proof.

The above result provides a condition for the effect of changing *C_k_*, the probability for transition from *A_k_*, on the stable population mean fitness. Specifically, if *A_k_* individuals have below-average fitness, then increasing *C_k_* will increase the population mean fitness.

We turn our attention to the case where the transition rates from a subset *k* of the types are correlated, that is, *C_j_* = *C_i_*, for *i, j* ∊ *k* In this case, Eq. 13 leads directly to the following.

### Corollary 1

*The sensitivity of the stable population mean fitness to change in the rate of transition τ from types in k is*

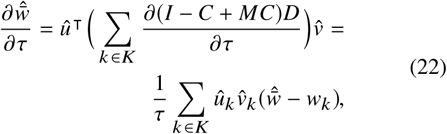

and

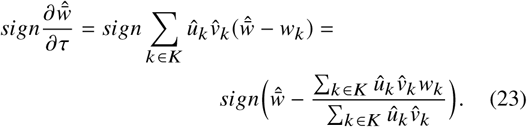

Therefore, increased transition from types in *k* will increase the stable population mean fitness if the average fitness of individuals descended from types in *k* is below the stable population mean fitness. For example, Ram and Hadany (2012, Appendix B) considered individuals that are grouped by the number of their accumulated mutant alleles, *k* (see Mutation model 2), and the effect of increasing the mutation rate in individuals with at least *π* mutant alleles. According to Eq. 23, this will result in increased stable population mean fitness if individuals with *π* or more mutant alleles have below-average fitness.

#### Reproductive value principle

An interesting interpretation of Eq. 16 is

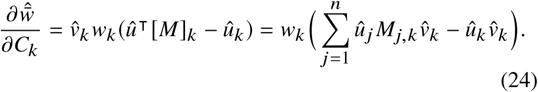

Here, *û_k_* can be regarded as the *reproductive value* of type *k* (Fisher, 1930, pg. 27), which gives the relative contribution of type *k* to the long-term population (see Appendix B). Consequently, 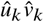 is the *ancestor frequency* of type *k* (Hermisson et al., 2002), namely the fraction of the equilibrium population descended from type *k*. The sum 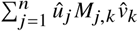 can be similarly interpreted as the fraction of the equilibrium population descended from individuals that transitioned from type *k* to another type (via the *k* column of the transition matrix *M*), conditioned on transition occurring.

Since *w_k_* > 0, from Eq. 16 we have the following corollary.

### Corollary 2 (Reproductive value principle)

*In the notation of Result 1,*

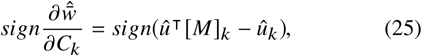

*where* [*M*]_*k*_ *is the k-th column of M*.

*Therefore, increased transition from type k will increase the stable population mean fitness if the fraction of the population descended from type k is expected to increase due to a transition to another type.*

Corollary 2 sheds light on why we require *M* to be primitive. If *M* is primitive then individuals of type *k* can transition into any other type in a finite number of generations. So individuals with below-average fitness can have descendants with above-average fitness, and increased generation of variation in these individuals will increase the stable population mean fitness. In contrast, if *M* is not primitive, individuals with below-average fitness are “doomed” and increasing the generation of variation in these individuals can only hasten their removal from the population. For example, if we set β = 0 in Mutation model 2, *M* becomes triangular and imprimitive, and the stable mean fitness becomes *w̅* = (1 – *δC*_0_)*w_0_*, which is not affected by changes in *C_k_* for *k* ≥ 1 (see also Agrawal, 2002; Ram and Hadany, 2012, Fig. 1A).

#### Evolutionary genetic stability

We now focus on a neutral modifier locus completely linked to the types *A_k_*, with no direct effect on fitness, and whose sole function is to determine *C_k_*, the rates of transition from the different types. We will show that modifier alleles that increase the stable population mean fitness in accordance with Result 1 are favored by natural selection.

#### Modifier model

Consider the case of two modifier alleles, *m* and *M*, inducing different transition probabilities *C* = diag [*C*_1_,…, *C_n_*] and 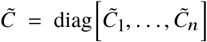, respectively. The frequencies of type *A_k_* linked to modifier *m* or *M* are *f_k_* and *g_k_*, respectively, where 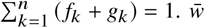 now ensures that 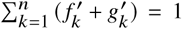, and the rest of the model parameters are the same as in Eq. 2.

The frequencies in the next generation, *f′* for allele *m* and *g′* for allele *M*, are given by

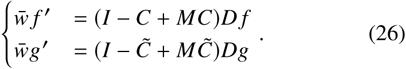

Here, 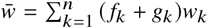 is the mean fitness of the entire population. Note that Eq. 2 is the special case of Eq. 26 where allele *M* is absent, i.e. *gk* = 0 for all *k*.

Result 11 provides a condition under which increasing the transition rate *C_k_* from type *A_k_* will increase the stable population mean fitness. Could a modifier allele that increases *C_k_* increase in frequency when initially rare in the population? To answer this we analyze the stability of resident modifier allele *m* with transition rates *C_k_* to invasion by a modifier allele *M* with rates 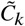 under Eqs. 26.

The equilibrium of Eqs. 26 when modifier allele *M* is absent from the population is 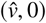, where 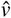 is given in Eq. 12 and *g_k_* = 0 for all *k*. The stability of allele *m* to invasion by allele *M* is determined by the leading eigenvalue *λ*1 of **L**_*ex*_ the external stability matrix of the equilibrium 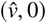, which, in turn, is determined by the Jacobian **J** of the system in Eqs. 26 evaluated at the equilibrium 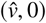, where

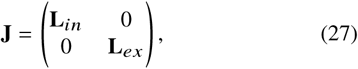

and **L**_*in*_ is the local stability matrix of the equilibrium 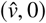 in the space 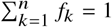. The zero block matrices are due to the complete linkage between the modifier and the types *A_k_* and to the lack of transition (i.e., mutation) between the modifier alleles.

**L**_*ex*_ can be written as

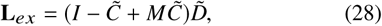

where 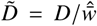 is the diagonal matrix with entries 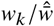 for all *k*, and 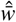 is the stable population mean fitness in the absence of the modifier allele *M*. *λ*_1_, the leading eigenvalue of **L**_*ex*_, coincides with 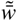, the maximal mean fitness associated with Eq. 28. Thus we can apply Result 1 to *λ*_1_ and obtain the following result.

##### Result 2 (Evolution of increased genetic variation)

*Let λ_1_ be the the leading eigenvalue of the external stability matrix* **L**_*ex*_. *If the transition rates induced by the modifier alleles m and M are equal, i.e.,* 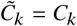 *for all k, then*

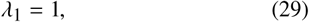

and

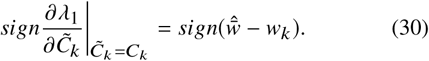

*Therefore, an initially rare modifier allele M with transition rates slightly different from the resident allele m can successfully invade the population (λ_1_* > 1*) if M increases the probability of transition from types with below-average fitness, thereby increasing the mean fitness* 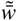.

###### Proof

Substituting 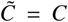 in Eq. 28 and multiplying both sides by 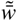,

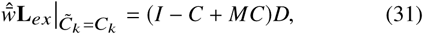

and since 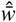 is the leading eigenvalue of the RHS (see Eq. 12), the leading eigenvalue of 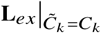 is *λ*_1_ = 1.

Now, applying Result 1 (Eq. 14) to Eq. 28, the sign of the derivative of *λ*_1_ with respect to 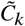 is

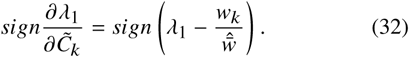

Thus

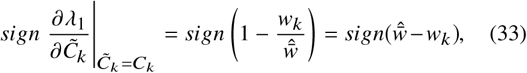

since 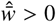. This completes the proof.

##### Reduction principle and mutational loss

Note that if the modifier has the same effect on all types, then we can substitute *C* = *μI* (with *μ* > 0) in Eq. 12, and proceeding as in Eq. 22, we find a relationship previously described by Hermisson et al. (2002, Eq. 24),

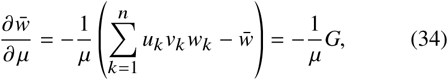

where *u_k_, V_k_* are computed at the equilibrium associated with *μ* and *w̅* is the mean fitness at that equilibrium. *G* is the difference between the *ancestral mean fitness* 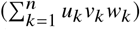 and the *stable population mean fitness* (*w̅*) when *C_k_* = *μ*, is called the *mutational loss* (Hermisson et al., 2002).

If an invading allele *M* changes the transition probability from that of the resident allele *m*, i.e., *C_k_* = *μ* and 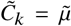, then **L**_*ex*_ becomes

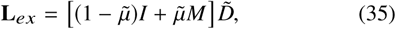

where 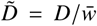 is the diagonal matrix with entries 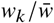 for all *k*. Using the *unified reduction principle* (Altenberg et al., 2017) the leading eigenvalue 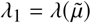 of **L**_*ex*_ satisfies 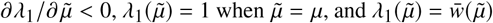. Thus we can conclude that the mutational loss *G* is positive, the reduction principle holds, and the mean fitness is a decreasing function of 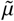.

## Discussion

We have shown that under constant-viability selection and in an effectively infinite haploid population at mutation-selection or migration-selection equilibrium, the stable population mean fitness increases if individuals with below-average fitness increase the rate at which variation is generated. Furthermore, modifier alleles that increase generation of variation in such individuals are favored by natural selection. These results apply as long as there is a chance for the variation-generating process to transform an individual with below-average fitness into one with above-average fitness (e.g. *M* in Eq. 2 is primitive).

We have given several examples of variation-generating processes for which this principle applies – namely mutation, migration, and learning (see *Models* section) – but our model may apply to other processes as well. For example, the *reduction principle* applies to ecological models of dispersal, and Gueijman et al. (2013) have demonstrated that even in homogeneous environments, fitness-associated dispersal increases the mean fitness of diploid populations and is favored by selection over uniform dispersal. Similarly, if the transmission fidelity of culturally-transmitted traits depends on the type or fitness of the transmitting individual, we expect that our results will hold (see Learning model).

Eq. 13 is a generalization of a result of Ram and Hadany (2012, Eq. 4). Ram and Hadany modeled the accumulation of mutant alleles in a population (see Mutation model 2). Using Eq. 36 in Appendix A and a recursion on the ratios of the reproductive values (see Ram and Hadany, 2012, eqs. A5-6), they concluded that at the mutation-selection balance, if individuals with below-average fitness 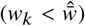 increase their mutation rate, then the population mean fitness will increase – a result generalized by our *Mean fitness principle* in Eqs. 14 and 23.

Our analysis focuses on populations at equilibrium. Nevertheless, it has been demonstrated that during adaptive evolution (i.e., in non-equilibrium populations), a modifier that increases the mutation rate of maladapted individuals can be favored by selection (Ram and Hadany, 2012; Lukačišinová et al., 2017) and increase the adaptation rate (Ram and Hadany, 2014), and empirical evidence suggests that *stress-induced mutagenesis* is common in bacteria and yeast, and may be prevalent in plants, flies, and human cancer cells (Rosenberg et al., 2012; Fitzgerald et al., 2017). Similar theoretical results have been demonstrated for a modifier that increases the recombination rate in maladapted individuals (Hadany and Beker, 2003a, b).

## Conclusions

Departures from the *reduction principle* for mutation, recombination, and migration rates usually involve fluctuating selection, non-equilibrium dynamics, or departures from random mating (see Carja et al. (2014) and references therein). Here we have provided another general example, which suggests that a modifier allele that causes individuals with below-average fitness to increase the rate at which variation is generated, will be favored by selection and will lead to increased population mean fitness.

## Appendices

### Appendix A

Caswell (1978) gave a *formula for the sensitivity of the population growth rate to changes in life history parameters*. In this formula, the *population growth rate* is the leading eigenvalue of the population transformation matrix T, the *life history parameters* are entries of *T*, and the *sensitivity* is the derivative of the former with respect to the latter. This is a useful formula (Caswell, 1978; Hermisson et al., 2002; Ram and Hadany, 2012; Otto and Day, 2007, ch. 10), and therefore we reproduce it here.

#### Lemma 1

*T be a non-negative matrix with leading eigenvalue λ and left aid right eigenvectors û and 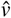 such that 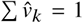 and 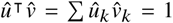. Then the sensitivity of λ to changes in any element t of the matrix T is*

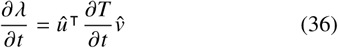

##### Proof

Using the lemma assumptions, 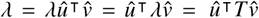 and differentiating both sides we get 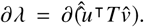 Using the product rule (once in each direction),

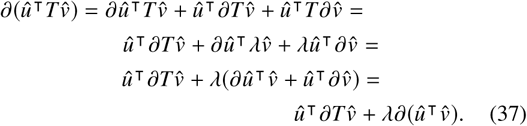

Because 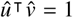, we have 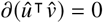 and 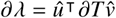.

### Appendix B

#### Remark 1 (Fisher’s reproductive value)

*Let M be an irreducible column-stochastic matrix and D be a positive diagonal matrix. The entries of the left Perron eigenvector û of the matrix MD can be regarded as* Fisher’s reproductive values *(Fisher, 1930, pg. 27)*

*Fisher’s reproductive values* can be understood as follows (Grafen, 2006; Otto and Day, 2007, ch. 10). Consider the dynamics not of frequencies but of absolute population sizes such that the vector of the number of individuals of each type at time *t* is *n*(*t*) and the corresponding frequencies are *f_k_* (*t*) = *n_k_* (*t*)/Σ_*i*_ *n_n_* (*t*). The dynamics are

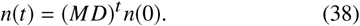

Let *n*(*k, t*) be the vector when the initial population is a single individual of type *k*. The dynamics are

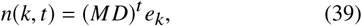

where *e_k_* is a vector with 1 at position *k* and 0 elsewhere.

The total population size at time *t* starting with type *k* is then

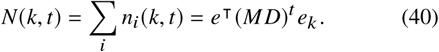

Now we can compare the sizes of populations based on what type they started with:

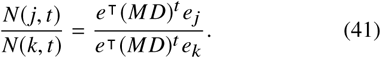

Now write *MD* in its Jordan canonical form

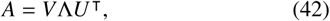

where *V* is the matrix of right (column) eigenvectors of *M D*, *U^T^* is the transposed matrix of left (row) eigenvectors of *M D*, where we can take *VU^T^* = *U^T^V* = *I*, and Λ is the diagonal matrix of eigenvalues of *A* (for a non-generic set of matrices *M*, the geometric and algebraic multiplicities of the eigenvalues of *M D* differ, and Λ will not be a diagonal matrix, a case we can ignore).

Hence,

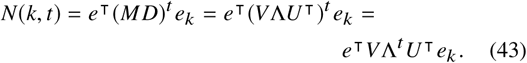

For the ratio, we can divide Λ by *λ*_1_ = *ρ*(*MD*), the spectral radius of *MD*:

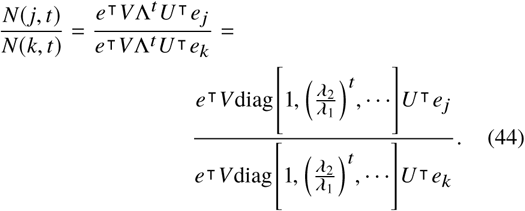

Now take the limit *t* → ∞. By assumption, *M D* is irreducible, so *λ_i_* < *λ*_1_ for all *i* > 1. Therefore,

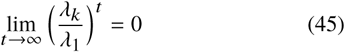

for all *k* > 1, and

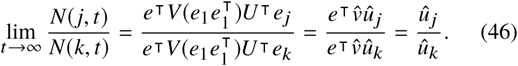

The vector *û* is the left *Perron* eigenvector of *M D*, and *û_k_* is *k*-th element of *û*. This is why the value *û_k_* can be interpreted as the *reproductive value* of type *k*: it is a weighting for the size of the population generated by a single individual of type *k*.

If we begin with a population at the equilibrium distribution 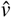, and ask what fraction of long-term descendants descended from type *k* at that time, we weight the equilibrium frequency 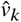 by the reproductive value 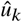, to get 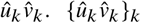 is a probability distribution, since

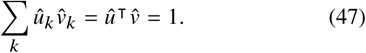

Hermisson et al. (2002) called this distribution the *ancestor* or *ancestral distribution.*

### Appendix C

#### Lemma 2 (Stability of the equilibrium 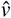)

*If C and D are positive diagonal matrices, and M is primitive, then the equilibrium 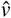 of the system in Eq. 12 is globally stable.*

##### Proof

Let *A* = (*I* – *C* + *MC*)*D* and denote the leading eigenvalue of A as 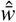.

According to the *Perron-Frobenius theorem,*

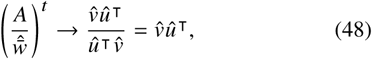

where *û* and 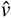 are the left and right Perron eigenvectors of *A* such that

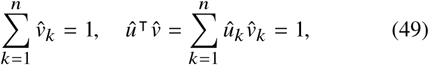

and 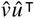 is the *Perron projection* into the eigenspace corresponding to 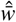.

Therefore, for any positive frequency vector *f*

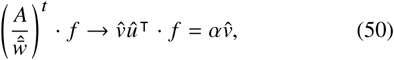

where *α* ∊ ℝ. Using eqs. 49 and 50

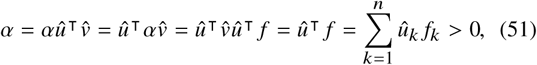

because *f* and *û* are positive vectors.

Now, 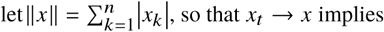

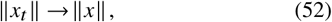

as *t* → ∞ and rewrite Eq. 2 as

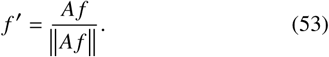

It is easily seen that *f^t^* = *A^t^f*/║*A^t^f*║. Hence

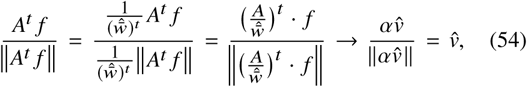

since 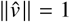 (Eq. 49) and *α* ≠ 0 (Eq. 51). This completes the proof.

## Acknowledgements

This work was supported in part by the Department of Information and Computer Sciences at the University of Hawai‘i at Mānoa, the Konrad Lorenz Institute for Evolution and Cognition Research, the Mathematical Biosciences Institute through National Science Foundation Award #DMS 0931642, the Stanford Center for Computational, Evolutionary and Human Genomics, and the Morrison Institute for Population and Resources Studies, Stanford University.

